# Comparative transcriptional profiling of the early host response to infection by typhoidal and non-typhoidal *Salmonella* serovars in human intestinal organoids

**DOI:** 10.1101/2020.11.25.397620

**Authors:** Basel H. Abuaita, Anna-Lisa E. Lawrence, Ryan P. Berger, David R. Hill, Sha Huang, Veda K. Yadagiri, Brooke Bons, Courtney Fields, Christiane E. Wobus, Jason R. Spence, Vincent B. Young, Mary X. O’Riordan

## Abstract

*Salmonella enterica* represents over 2500 serovars associated with a wide-ranging spectrum of disease; from self-limiting gastroenteritis to invasive infection caused by non-typhoidal serovars (NTS) and typhoidal serovars, respectively. Host factors strongly influence infection outcome as malnourished or immunocompromised individuals can develop invasive infections from NTS, however, comparative host responses to individual serovars have been difficult to perform due to reliance on poorly representative model systems. Here we used human intestinal organoids (HIOs), a three-dimensional “gut-like” *in vitro* system derived from human embryonic stem cells, to elucidate similarities and differences in host responses to NTS and typhoidal serovars. HIOs discriminated between the two most prevalent NTS, *Salmonella* enterica serovar Typhimurium (STM) and *Salmonella* enterica serovar Enteritidis (SE), and typhoidal serovar *Salmonella* enterica serovar Typhi (ST) in epithelial cell invasion, replication and transcriptional responses. Pro-inflammatory signaling and cytokine output was reduced in ST-infected HIOs compared to NTS infections, reflecting early stages of NTS and typhoidal diseases. While we predicted that ST would induce a distinct transcriptional profile from the NTS strains, more nuanced expression profiles emerged. Notably, pathways involved in cell cycle, metabolism and mitochondrial functions were downregulated in STM-infected HIOs and upregulated in SE-infected HIOs. These results correlated with elevated levels of reactive oxygen species production in SE-infected HIOs compared to mock-infected HIOs. Collectively, these results suggest that the HIO model is well suited to reveal host transcriptional programming specific to individual *Salmonella* serovars, and that individual NTS may provoke unique host epithelial responses during intestinal stages of infection.

**Author Summary:** *Salmonella enterica* is the major causative agent of bacterial infections associated with contaminated food and water. *Salmonella enterica* consists of over 2500 serovars of which Typhimurium (STM), Enteritidis (SE) and Typhi (ST) are the three major serovars with medical relevance to humans. These serovars elicit distinctive immune responses and cause different diseases in humans, including self-limiting diarrhea, gastroenteritis and typhoid fever. Differences in the human host response to these serovars are likely to be a major contributing factor to distinct disease outcomes but are not well characterized, possibly due to the limitations of human-derived physiological infection models. Unlike immortalized epithelial cell culture models, human intestinal organoids (HIOs) are three-dimensional structures derived from embryonic stem cells that differentiate into intestinal mesenchymal and epithelial cells, mirroring key organizational aspects of the intestine. In this study, we used HIOs to monitor transcriptional changes during early stages of STM, SE and ST infection. Our comparative analysis showed that HIO inflammatory responses are the dominant response in all infections, but ST infection induces the weakest upregulation of inflammatory mediators relative to the other serovars. In addition, we identified several cellular processes, including cell cycle and mitochondrial functions, that were inversely regulated between STM and SE infection despite these serovars causing similar localized intestinal infection in humans. Our findings reinforce HIOs as an emerging model system to study *Salmonella* serovar infection, and provide global host transcriptional response profiles as a foundation for understanding human infection outcomes.

## Introduction

*Salmonella enterica* greatly impacts human health causing an estimated 115 million infections worldwide every year and are one of the four leading causes of diarrheal diseases [1,2]. *Salmonella enterica* is a copious species consisting of over 2500 serovars and infects the intestinal epithelial layer causing diseases ranging from asymptomatic carriers to more severe systemic disease. *Salmonella* serovars are classified based on host specificity and disease outcomes. Host generalist serovars including *Salmonella enterica* serovar Typhimurium (STM) and Enteritidis (SE) infect a broad range of hosts and cause localized inflammation and self-limiting diarrhea in healthy individuals or more severe gastroenteritis in children and elderly. In contrast, host-restricted serovars including Typhi and Abortusovis infect only one host and cause more serious clinical manifestations including Typhoid fever in humans and abortions in mares respectively.

Although *Salmonella enterica* serovars share a conserved core genome, determinants of host specificity and varying clinical manifestations are poorly understood. The molecular basis for distinct host adaptation and disease outcome is likely to be multifactorial, mediated by bacteria and host-dependent mechanisms. Initial comparative genomic analyses identified specific signatures that may be indicative of some of these differences [3]. However, comparative host signatures across different serovars are still limited by host specificity and poorly representative model systems. Using human epithelial cell lines addresses host-specificity, but immortalized cell lines do not represent the multiple subsets of intestinal epithelial cells found in the gut and harbor mutations that likely alter cellular responses to bacterial infection.

Human intestinal organoids (HIOs) have emerged as an alternative *in vitro* model to study intestinal epithelial host responses to commensal microbiota and enteric pathogens. HIOs are differentiated from pluripotent stem cells into three-dimensional spheroids composed of a defined luminal space bound by a polarized epithelial barrier surrounded by mesenchyme. This is an improvement over existing models because the untransformed HIO epithelium is polarized, and contains multiple epithelial cell lineages found in the intestine [4]. Hill *et al*. showed that HIOs supported luminal growth of *Escherichia coli* following microinjection, and that physiological changes in the HIO occurred during colonization, such as an increase in mucus production, mirroring what happens *in vivo* during initial colonization [5]. Forbester *et al*. also showed that STM invades HIO epithelial cells and induces inflammatory responses. Here, we used HIOs to compare the transcriptome of intestinal epithelial responses to host-restricted *Salmonella enterica* serovar Typhi and two host unrestricted serovars Typhimurium and Enteritidis. We found that *Salmonella* infection induced a variation in magnitude of immune responses that was dependent on the infecting serovar. ST infection induced the weakest response, which is consistent with the idea that ST actively suppresses host responses to establish a systemic infection. Notably, we found that both STM and ST infection similarly decreased expression of pathways involved in cell cycle, DNA repair and DNA replication while SE infection increased these responses.

## Results

### *Salmonella* serovars invade HIO epithelial cells and induce distinct patterns of mucus production

To study initial host responses to *Salmonella*, we micro-injected bacteria into the luminal space of the HIO to allow luminal replication throughout the course of infection. This model better resembles the continuous interaction between bacteria and intestinal cells during the natural course of infection. We first determined whether different *Salmonella* serovars could colonize and invade HIO epithelial cells. We chose the most prevalent serovars that cause gastroenteritis, STM and SE, and a typhoidal serovar, ST. HIOs were microinjected with STM, SE or ST, and total bacterial burden per HIO was enumerated at 2.5 and 24 hours post infection (h pi) **(Fig 1A)**. All serovars showed at least a 1.5 log increase in bacterial burden at 24h pi, relative to 2.5h pi. Intracellular bacterial burden was quantified by gentamicin protection assay, where HIOs were sliced open to expose luminal bacteria to gentamicin prior to quantifying colony forming units **(Fig 1B)**. Intracellular bacterial numbers increased over time with all serovars, suggesting that intracellular replication or continued invasion contributes to the increase in bacterial load at 24h pi. Noticeably, STM infection resulted in a slightly higher bacterial burden in both total and intracellular CFU at 24h pi, compared to the other serovars. This observation suggests that STM invades HIO epithelial cells more efficiently than SE and ST. We also evaluated epithelial morphology by performing hematoxylin and eosin (H&E) staining **(Fig 1C).** HIOs remained intact during infection with all serovars for the duration of the experiment. However, damaged regions of the HIO epithelial lining could sometimes be observed, especially during STM infection. In addition, periodic acid-schiff reagent (PAS) and alcian blue (AB) staining was also performed to detect mucus, as a recent study showed that HIOs increase mucin production during bacterial colonization [5]. In agreement with these findings, PAS and AB staining revealed an increase in mucus production in response to infection with all three serovars **(Fig 1D)**. Interestingly, we observed unique staining patterns during infection with the different serovars. In ST-infected HIOs, mucus appeared to accumulate within cells, while STM and SE infection resulted in evacuation of mucus into the HIO lumen. Thus, within a 24h period, all three *Salmonella* serovars can colonize and invade HIOs, inducing distinct patterns of mucus production without causing major destruction to the HIO epithelial layer.

**Fig 1.**
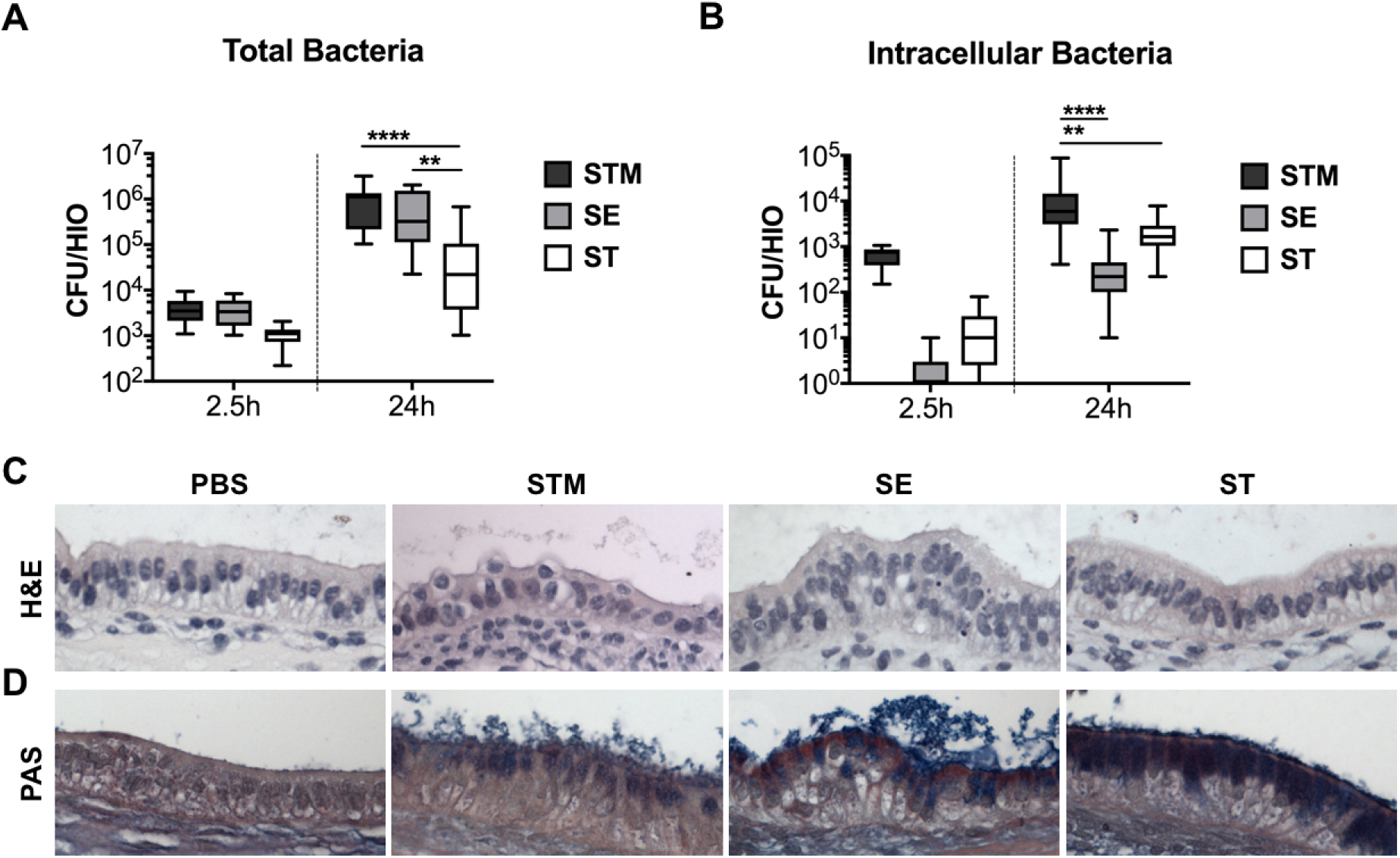
*Salmonella enterica* serovars invade HIO epithelial cells and stimulate mucin production. (A) Total bacterial burden was enumerated per HIO at 2.5h and 24h post injection. (B) Intracellular bacterial burden was enumerated after exposing luminal bacteria to gentamicin at 2.5h and 24h post injection. (C and D) Histology sections of HIOs at 8h pi using Hematoxylin and Eosin (H&E) staining (C) and Periodic acid-Schiff (PAS) staining (D). Whiskers represent min and max values of n > 16 HIOs from three independent experiments. Statistical significance within the group was determined by two-way ANOVA and followed up by Tukey’s multiple comparisons test. *P* values < 0.05 were considered significant and designated by: \*\**P* < 0.01 and **** *P* < 0.0001.

### Host transcriptional dynamics differ between *Salmonella* serovars

To define the global host transcriptional response to the 3 *Salmonella* serovars, we performed RNA sequencing (RNA-seq) at 2.5h and 8h pi with HIOs that were infected with STM, SE or ST and compared transcript levels to control PBS-injected HIOs. Principal component analysis (PCA) was performed on normalized gene counts to identify clustering patterns between conditions **(Fig 2A)**. The PCA plot showed clear segregation and clustering of samples based on both infection and time. Infected HIOs at 2.5h had the most variation relative to PBS-injected HIOs where they were separated by the first (the greatest variance) principal component and further clustered based on infection with each serovar. STM-infected HIOs showed the greatest separation from the control, while SE-infected HIOs showed an intermediate and ST-infected HIOs showed the least separation. To further investigate HIO responses during *Salmonella* infection, we identified differentially expressed genes (DEGs) between HIOs injected with PBS and infected with STM, SE or ST **(S1 and S2 Tables)**. We found that there were comparable numbers of DEGs during infection of all serovars at 2.5h pi **(Fig 2B)**. Some of the DEGs were shared between infections with each serovar, which likely represents a core host response to *Salmonella* infection. However, infection with each serovar also resulted in induction and suppression of a unique set of DEGs. Notably, transcriptional dynamics showed that there was an increase in the number of DEGs at 8h pi in response to infection with the non-typhoidal serovars (NTS), STM and SE, while the number of DEGs during infection with ST decreased **(Fig 2B)**. Collectively, the HIO responses represent two patterns; core transcriptional responses that are changed during *Salmonella* infection and serovar-specific responses.

**Fig 2.**
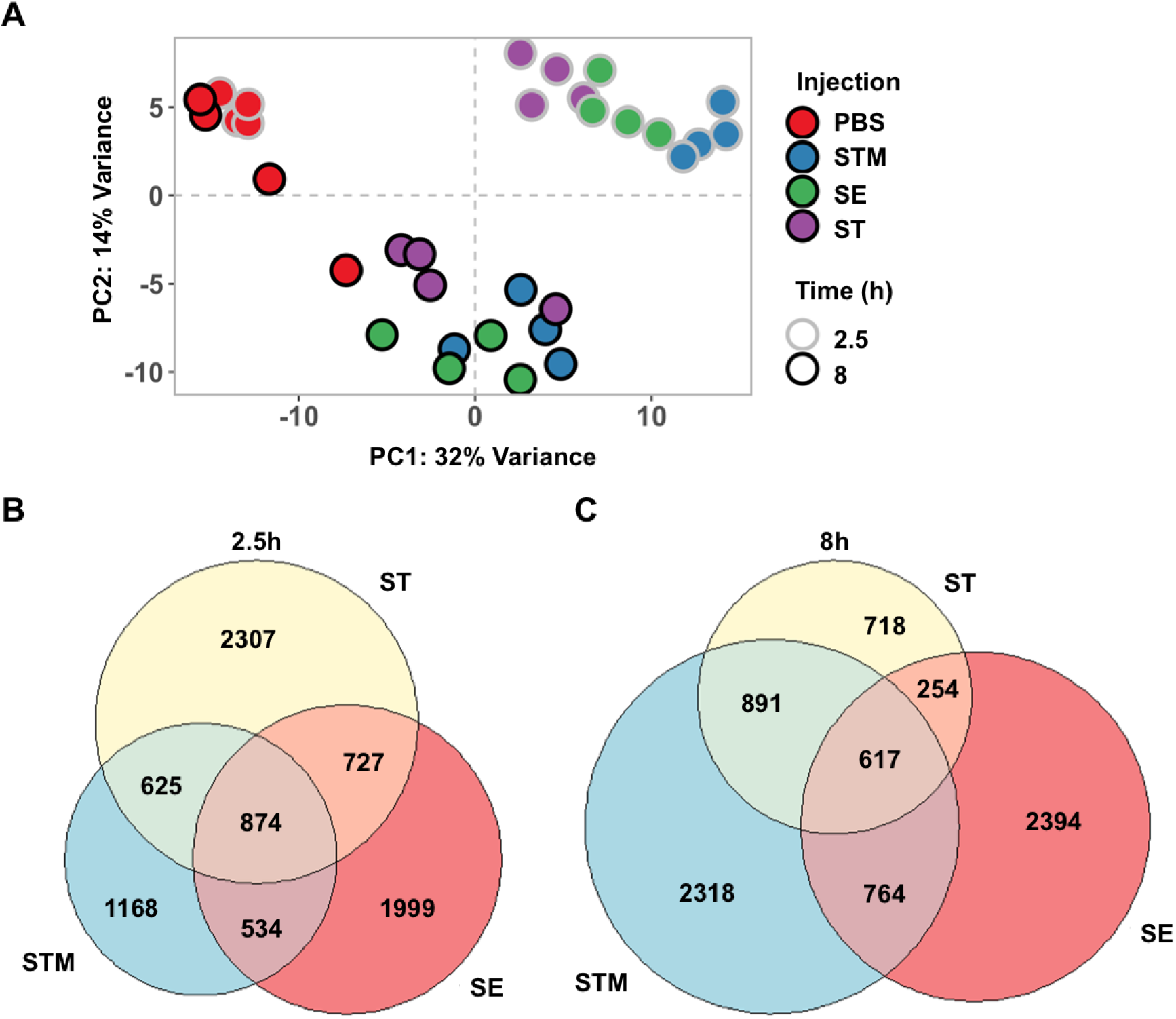
Changes in HIO gene expression is driven by both serovar and time post infection. (A) Principal component analysis of HIOs injected with various *Salmonella* serovars. Each circle represents a biological replicate of a pool of five HIOs. (B and C) Euler diagram comparison of gene expression changes in each HIO condition relative to PBS-injected HIOs at 2.5h (B) and 8h (C) post injection. Genes were filtered by *P* value < 0.05.

### *Salmonella* serovars differentially alter inflammatory, stress response, metabolism of lipid and amino acids and cell cycle pathways

To identify biological pathways associated with DEGs from each infection condition, gene sets were separated into upregulated (increased) and downregulated (decreased) categories based on fold change relative to PBS-injected HIOs and imported separately into the Reactome pathway analysis tool **(S3-S6 Tables)**. In the upregulated datasets, the majority of the significant pathways in all three infection conditions at both 2.5h and 8h belonged to the immune system and signal transduction category **(Fig 3A)**. We found that infection induced a complex response in both innate immune and cytokine signaling pathways including, but not limited to, Toll-like receptors, interleukin mediators and Type I interferons **(Fig 3B)**. Notably, only in ST-infected HIOs were some immune system pathways associated with downregulated DEGs, such as non-canonical NF-κB and Interleukin-1 signaling. These results revealed that inflammatory pathways were the primary responses during *Salmonella* infection and are consistent with the hypothesis that typhoidal serovar infection is relatively “silent”, producing less inflammatory mediators compared to NTS infection.

**Fig 3.**
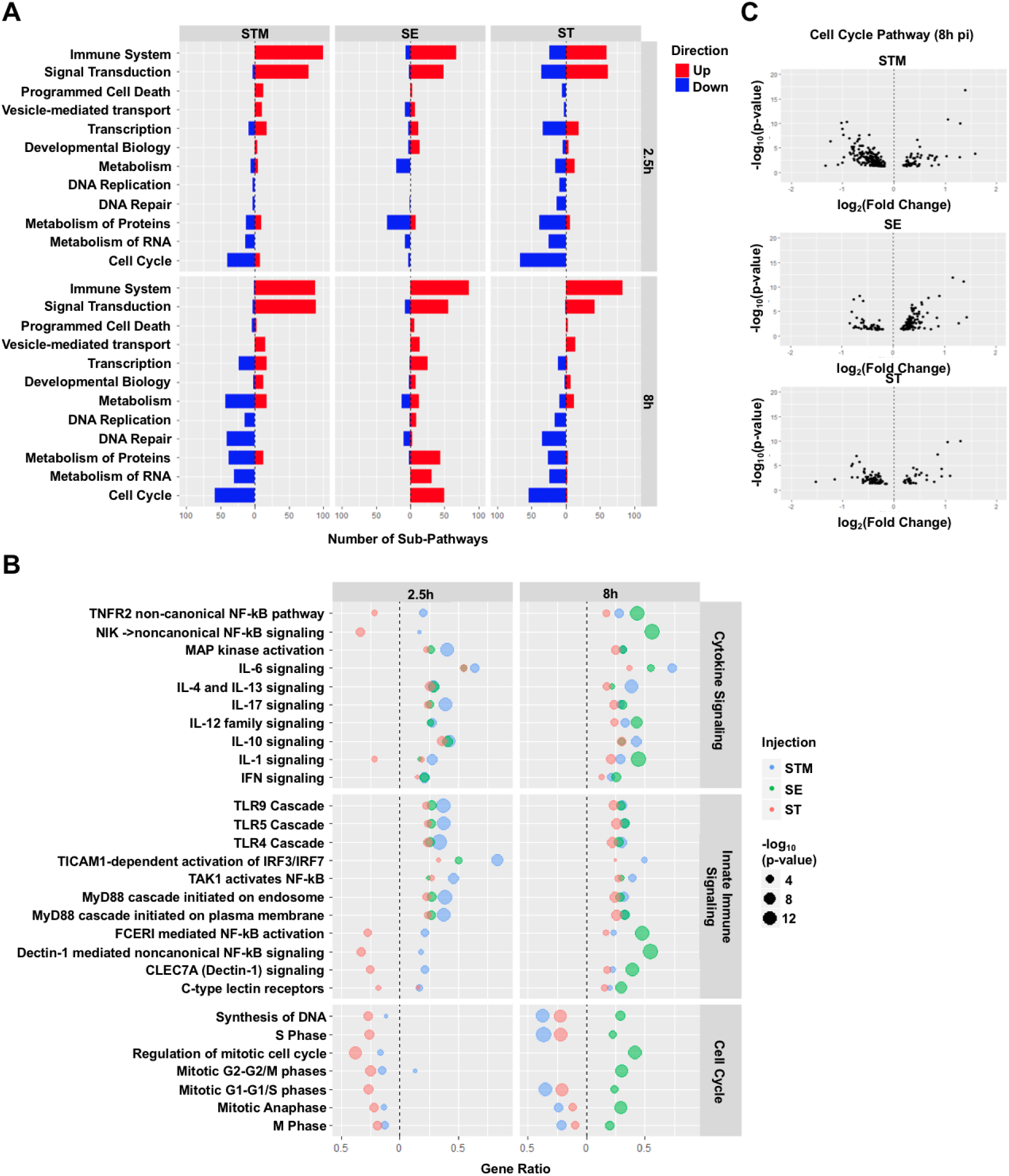
Immune system and cell cycle pathways encompass the predominant increases and decreases in gene expression during infection. (A) Number of sub-pathways clustering into hierarchy major Reactome cellular processes. Significantly upregulated (red) or downregulated (blue) genes were analyzed using ReactomePA and pathways were clustered into hierarchy cellular processes from the Reactome database. Hierarchy cellular processes with at least 12 significant sub-pathways in at least one infection condition were included. (B) Dot plot showing top pathways enriched from the Reactome database. Pathway coverage shown as gene ratio with significantly upregulated pathways shown on the right of the dotted line and downregulated pathways on the left. Dot size represents–log_10_(p-value) of enriched pathway during HIOs infection with STM in blue, SE in green or ST in red. (C) Volcano plots of cell cycle genes that changed significantly during infection with STM, SE or ST. Significant genes and pathways were determined based on *P* value < 0.05.

Apart from the predominant inflammatory pathways, we also identified several differentially upregulated pathways in response to *Salmonella* serovars that have been linked to intestinal infection. These pathways included antigen presentation, extracellular matrix organization (ECM), lipid and amino acid metabolism and cellular stress responses including IRE1α-mediated unfolded protein response (UPR), mitophagy and the inflammasome (**S1 Fig**). Although there were genes in these pathways that were significantly upregulated in response to all three serovars, some were enriched only in response to a specific serovar. For example, we found that pathways belonging to ECM, UPR and tryptophan catabolism were significantly upregulated at 8h pi during STM infection but not during SE and ST infection. In contrast, we found that cholesterol metabolism pathways were highly enriched in ST infection while amino acid metabolism, cellular responses to hypoxia, the inflammasome and antigen presentation pathways were significantly induced only in SE-infected HIOs.

Most of the significantly down-regulated pathways during STM and ST infections belonged to cell cycle, DNA replication and repair, metabolism of protein and metabolism of RNA **(Fig 3A)**, which point to a potential reduction in cell proliferation. Interestingly, in SE-infected HIOs, some of these categories including cell cycle and metabolism of protein were instead associated with upregulated DEGs at 8h pi **(Fig 3A and B)**. To further investigate how cell cycle genes changed in response to each serovar, we generated volcano plots to identify the distribution of significant cell cycle genes in response to infection **(Fig 3C-E)**. The majority of the cell cycle DEGs were downregulated at 8h in response to both STM and ST infection despite there being fewer significant DEGs associated with ST infection. In contrast, most of the cell cycle DEGs were upregulated in response to SE infection, suggesting that STM and ST may reduce HIO cell proliferation while SE infection may uniquely increase it. Taken together, we found that while most inflammatory responses are commonly upregulated, other responses, including ECM, cellular stress responses, lipid and amino acid metabolism and cell cycle are differentially regulated upon infection with these three *Salmonella* serovars.

### *Salmonella* serovars induce distinct HIO proinflammatory response profiles

Intestinal epithelial cells initiate inflammatory responses via production of proinflammatory mediators. Because the most dramatic transcriptional responses we observed were related to immune signaling, we sought to identify the HIO signature of inflammatory mediators including chemokines, cytokines and antimicrobial peptides (AMP) in response to each *Salmonella* serovar **(Fig 4A and S2 Fig)**. We found that all of these mediators were induced early during infection although with different magnitudes. For example, Colony Stimulating Factor 3 (CSF3), Interleukin 17C (IL17C), Interleukin 19 (IL19), C-C Motif Chemokine Ligand 20 (CCL20), C-X-C Motif Chemokine Ligand 1 (CXCL1), Defensin beta 4A (DEF4A) and Peptidase Inhibitor 3 (PI3) were highly induced during STM infection, moderately induced during SE infection but only weakly induced during ST infection. Of interest, IL17C signaling regulates epithelial host defense against mouse enteric pathogens [6]. HIO production of IL17C and its known downstream proinflammatory mediators, including CSF3 and DEF4A, also suggest that IL17C signaling modulates human intestinal host defense against *Salmonella* infection. To validate these transcriptional results, we measured production of specific inflammatory mediators (cytokine, chemokines and AMP) in the HIO culture medium by ELISA. All three serovars induced production of these mediators **(Fig 4B and S3 Fig)**. In general, changes in protein level correlated with transcriptional changes observed in our RNA-seq dataset, where STM-infection resulted in the highest levels of cytokine production, SE-infection resulted in an intermediate phenotype and ST-infection induced the lowest levels. Collectively, the data indicate that each serovar, even the two non-typhoidal serovars, interacts distinctly with the host to tune production of inflammatory mediators during infection. Our data also reflect previous reports that ST infection is less inflammatory than infection by other *Salmonella* serovars and suggest that HIOs are a useful platform for studying ST interactions with human epithelium.

**Fig 4.**
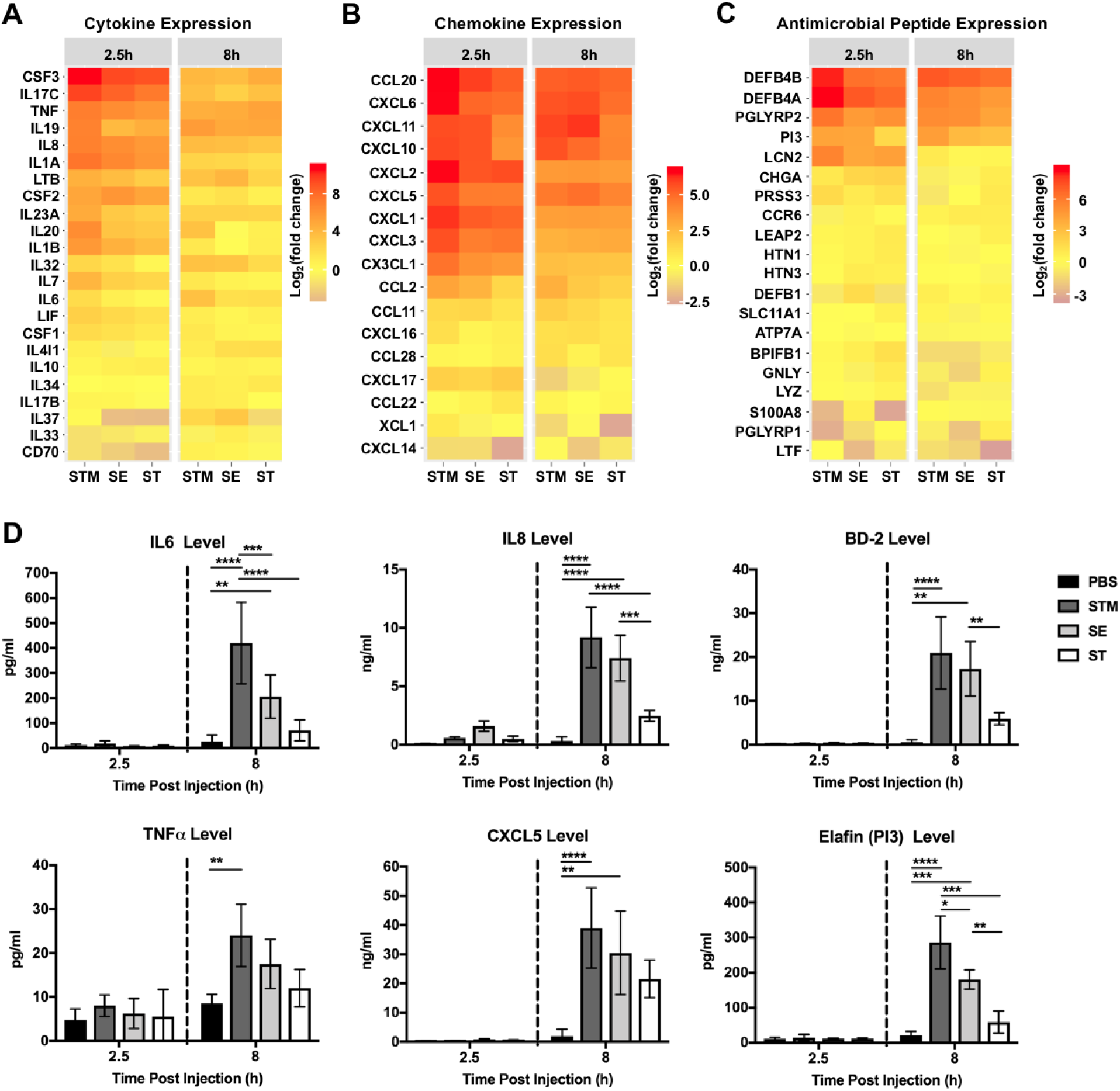
Differential gene expression and secretion of immune modulators by HIOs in response to infection. Gene expression of Cytokine (A), Chemokine (B) and Antimicrobial peptide (C) are presented as log_2_ fold change relative to PBS-injected HIOs at 2.5h and 8h post injection. (D) Cytokine, chemokine and antimicrobial peptide levels measured from HIO supernatant at 2.5h and 8h post injection via ELISA. n = 4 biological replicates. Error bars represent SD. Significance calculated by two-way ANOVA. *P* values < 0.05 were considered significant and designated by: \**P* < 0.05, \*\**P* < 0.01, \*\*\**P* < 0.001and **** *P* < 0.0001.

### Mitochondrial processes are differentially regulated during NTS infections

Although NTS cause similar disease manifestations in humans, they may interact with the intestinal epithelium by varied mechanisms as their genomes contain some different accessory genes [3]. Our data indicated that one of the most differentially regulated cellular processes between NTS was related to metabolism of proteins **(Fig 3A)**. To further identify major pathways within this category that were differentially regulated during infection with NTS, we sorted the significant pathways that belonged to the metabolism of proteins category in the Reactome database to identify these pathways.

We found that pathways belonged to three major categories; translation, protein folding, and post-transcriptional regulation that were increased in SE-infected HIOs but decreased during STM infection **(Fig 5A)**. Within the translation umbrella category, we found many mitochondrial-related processes, including mitochondrial translation, mitochondrial protein import and oxidative phosphorylation, were increased during SE infection but decreased during STM infection **(Fig 5B)**, suggesting that mitochondrial functions may differentiate between the host response to NTS during early stages of infection. Because mitochondria produce reactive oxygen species (ROS) during metabolism, we monitored generation of ROS in HIOs during infection **(Fig 5C)**. Consistent with an increase in mitochondrial gene expression, SE infection led to an accumulation of ROS in the HIOs when compared to PBS-injected HIOs. ROS induction was specific to SE, as STM infection did not trigger ROS generation when monitored at 24h pi. Collectively, our results suggest that SE infection induces specific HIO responses, including induction of mitochondrial related processes and ROS generation that distinguish this serovar from the more well-studied STM serovar.

**Fig 5.**
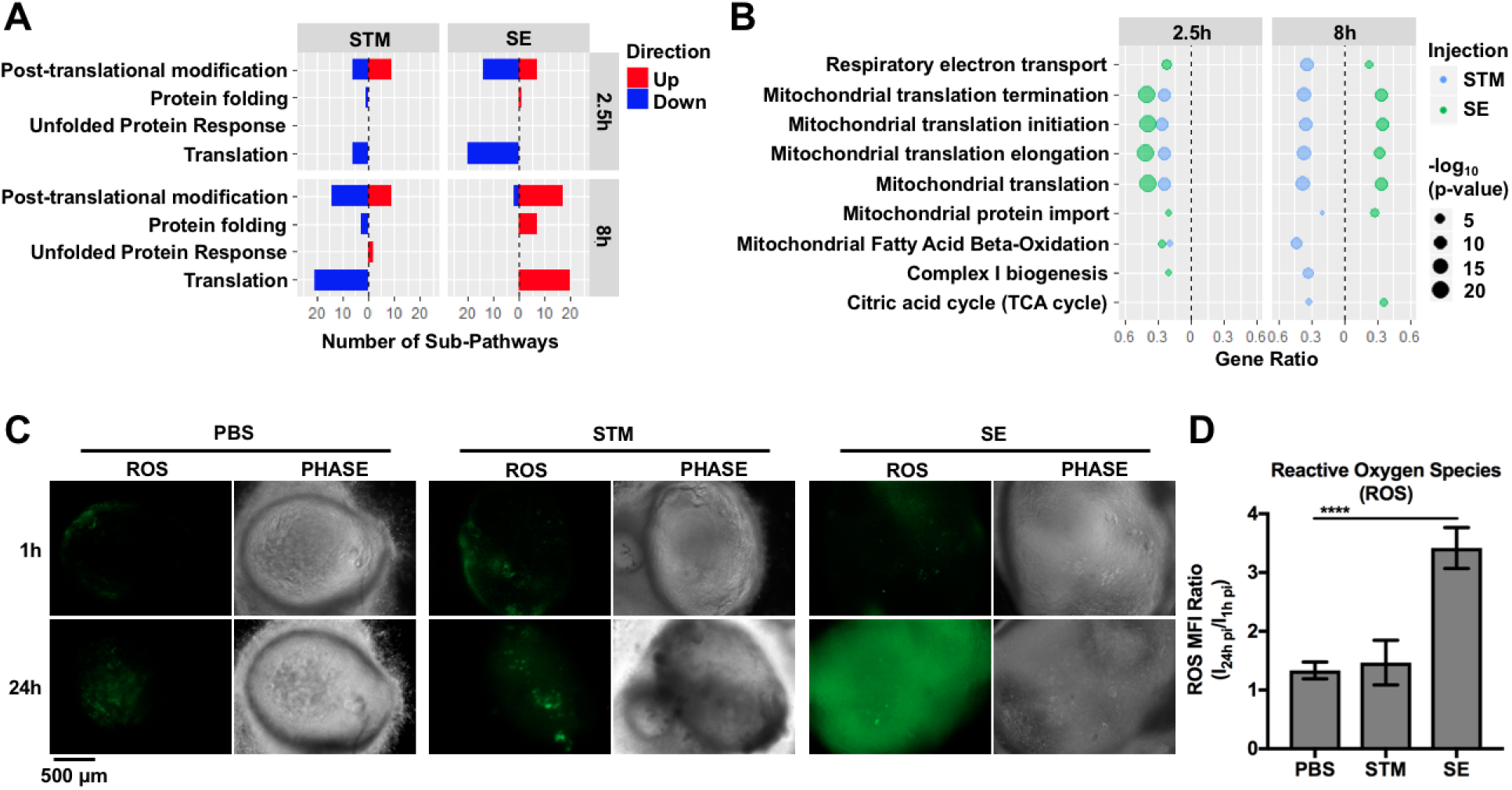
NTS infections inversely regulate changes in mitochondrial-related cellular processes at 8h pi and trigger differential ROS production. (A) Number of significant sub-pathways from the Reactome metabolism of proteins category during NTS infection. Upregulated (red) or downregulated (blue) pathways were identified using ReactomePA. Significant pathways were determined based on *P* value < 0.05. (B) Dot plot showing significantly enriched mitochondrial metabolism-related pathways in NTS-infected HIOs. Filtered by *P* value < 0.05. (C) Representative fluorescence images measuring reactive oxygen species levels in HIOs by general oxidative stress dye, CM-H2DCFDA. (D) Quantitation of reactive oxygen species levels of (C) with n ≥ 6 HIOs. Error bars represent SD. Statistical significance was determined by one-way ANOVA followed up by Tukey’s multiple comparisons test. **** *P* < 0.0001.

## Discussion

Despite sharing high genome identity, some *Salmonella* serovars cause infections that remain localized in the intestine while others cause more severe systemic infection. Differential host responses, especially the initial interactions with the intestinal epithelium, are likely a contributing factor in determining infection outcome, something that has been difficult to study with other established infection models. Here we describe the first comparative transcriptomic analysis using human intestinal organoids infected with *Salmonella* serovars Typhimurium, Enteritidis and Typhi. We compared HIO transcriptional profiles at different time points during infection and identified responses that were similar and unique to each serovar. As expected, inflammatory responses were a dominant early response during infection with all three serovars. However, at later times post infection, we observed distinct responses to each serovar including differences in expression of genes in cell cycle and mitochondrial function-related pathways. Direct comparison of HIO responses to these serovars revealed that many pathways that were decreased during STM infection were, in contrast, increased during SE-infection even though these serovars cause similar diseases in humans. Thus, our data highlight the utility of the HIO model to define signatures of host responses to closely related enteric pathogens to understand how these responses may shape disease manifestations.

ST is a human-specific pathogen and therefore it is most physiologically relevant to use human-derived cells to define mechanisms of pathogenesis and host response. Many previous studies of ST infection employed transformed cell lines [7,8], which may overlook particular aspects of epithelial function. In the HIOs, we observed that some inflammatory and signal transduction pathways were decreased early during ST infection when compared to NTS. However, this effect was abrogated by 8h pi, emphasizing that ST may specifically modulate intestinal epithelial responses early and transiently during infection. We also compared transcript and protein levels of immune mediators including cytokines, chemokines and defensins in response to ST and NTS. We demonstrated that ST infection produced less IL-6, IL-8, BD-2 and ELAFIN when compared to STM and SE infection despite comparable transcript levels. These findings lead us to propose that ST may block the ability of intestinal epithelial cells to release immune mediators post-transcriptionally. Several mechanisms have recently been shown to control cytokine and chemokine production post-transcriptionally [9–11]. Whether these mechanisms control ST pathogenesis and intestinal inflammatory responses are unknown, but could be elucidated in the HIO model.

Our finding that the three *Salmonella* serovars showed ifferential regulation of cell cycle pathways was intriguing. Intestinal epithelial cells undergo self-renewal to maintain barrier integrity, and infection with enteric pathogens can accelerate or inhibit cell proliferation to gain a survival advantage in the gut [12]. For example, *Citrobacter rodentium* stimulates the proliferation of undifferentiated epithelial cells, which increases oxygenation of the mucosal surface in the colon to create a replicative niche [13]. By contrast, some enteric pathogens including STM, *H. pylori* and *Shigella* are equipped with virulence factors to counteract intestinal cell proliferation and rapid epithelial turnover to enhance virulence [12]. In our experiments, both STM and ST infections resulted in downregulation of many genes in the cell cycle pathway while SE infection resulted in upregulation of several of these genes. Of note, it was previously reported that STM blocks epithelial cell proliferation via Type 3 Secretion System-2 effectors SseF and SseG [14]. These effectors are also encoded in the ST and SE genomes, but it is unclear whether expression levels or kinetics of SseF and SseG might differ to allow fine-tuned control of cell proliferation and pathogenesis.

Although STM and SE cause similar diseases in humans, we were surprised to observe that these two serovars exhibited the most variation in HIO responses relative to each other, including regulation of mitochondrial function-related genes. Our prior research has been focused on how cellular stress pathways contribute to innate immunity and we have recently shown that mitochondrial ROS contributes to bacterial killing by macrophages [15]. Interestingly, we observed that many pathways involved in mitochondrial metabolism are upregulated during SE infection and downregulated during STM infection. Accordingly, we found that SE infection increased generation of antimicrobial ROS in the HIOs, suggesting that an increase in mitochondrial metabolism may be important in intestinal host defense. Indeed, mitochondrial integrity and function is required for the maintenance of healthy intestinal barriers to prevent bacterial translocation across the epithelial lining [16,17]. In addition, recent studies demonstrated that metabolites produced by microbes in the gut can influence mitochondrial biogenesis and inflammation [18]. Given that both STM and SE are present in the HIO lumen through the course of infection, it remains unclear whether SE uniquely increases expression of mitochondrial genes, or luminal bacteria generally increase expression of mitochondrial genes but STM uniquely decreases their expression, or both. SE encodes more than 200 genes that are absent in either the STM or ST genome, which are clustered in unique islands designated as “regions of difference” (ROD) [19]. Some of these additional genes have been linked to SE pathogenesis using a mouse model of *Salmonella* infection [20,21]. Therefore, we speculate that genes expressed only by SE might account for SE-specific HIO responses and further work is required to elucidate mechanisms by which SE induces these specific responses.

Altogether, our findings show that the HIOs are a productive model to study early interactions of Salmonella serovars with the intestinal epithelium. HIOs have been previously used to probe for transcriptional responses during STM infection [22], but to our knowledge this is the first study to directly compare non-transformed human intestinal epithelial responses between non-typhoidal and typhoidal serovars. Looking beyond the pro-inflammatory pathways induced during infection by all three serovars, we identified unique host responses that are individually associated with these closely related serovars. Patterns emerging from our HIO experiments open up avenues for future studies to elucidate mechanisms by which different serovars fine-tune inflammatory output and modulate cell cycle and mitochondrial functions.

## Materials and Methods

### HIO Differentiation and Culture

HIO were generated by the I*n Vivo* Animal and Human Studies Core at the University of Michigan Center for Gastrointestinal Research as previously described [23]. Briefly, human ES cell line WA09 was obtained from Wicell International Stem Cell Bank and cultured on Matrigel-coated (BD Biosciences) 6-well plates in mTeSR1 media (Stem Cell Technologies) at 37°C in 5% CO2. Cells were seeded onto Matrigel-coated 24-well plates in fresh mTeSR1 media and grown until 85-90% confluence. Definitive endoderm differentiation was induced by washing the cells with PBS and culturing in endoderm differentiation media (RPMI 1640, 2%FBS, 2 mM L-glutamine, 100 ng/ml Activin A, 100 U/ml of Penicillin and 100 μg/ml of Streptomycin) for three days where fresh medium was added each day. Cells were then washed with endoderm differentiation media without Activin A and cultured in mid/hindgut differentiation media (RPMI 1640, 2%FBS, 2 mM L-glutamine, 500 ng/ml FGF4, 500 ng/ml WNT3A, 100 U/ml of Penicillin and 100 μg/ml of Streptomycin) for 4 days until spheroids were present. Spheroids were collected, mixed with ice cold Matrigel (50 spheroids + 50μl of Matrigel + 25μl of media), placed in the center of each well of a 24-well plate, and incubated at 37°C for 10 min to allow Matrigel to solidify. Matrigel embedded spheroids were grown in ENR media (DMEM:F12, 1X B27 supplement, 2 mM L-glutamine, 100 ng/ml EGF, 100 ng/ml Noggin, 500 ng/mL Rspondin1, and 15 mM HEPES) for 14 days where media were replaced every 4 days. Spheroids growing into organoids (HIOs) were dissociated from the Matrigel by pipetting using a cut wide-tip (2-3 mm). HIOs were mixed with Matrigel (6 HIOs + 25μL of media + 50μL of Matrigel) and placed in the center of each well of 24-well plates and incubated at 37°C for 10 min. HIOs were further grown for 14 days in ENR media with fresh media every 4 days. Prior to experiments, HIOs were carved out of the Matrigel, washed with DMEM:F12 media, and re-plated with 5 HIO/well in 50μL of Matrigel in ENR media with media exchanged every 2-3 days for 7 days prior to microinjection.

### Bacterial Growth Conditions and HIO Microinjection

*Salmonella* strains used in this study are *Salmonella* enterica serovar Typhimurium strain SL1344, *Salmonella* enterica serovar Enteritidis strain P125109 and *Salmonella* enterica serovar Typhi strain Ty2. Strains were stored at −70°C in LB medium containing 20% glycerol and cultured on Luria-Bertani (LB, Fisher) agar plates. Selected colonies were grown overnight at 37ºC under static conditions in LB liquid broth. Bacteria were pelleted, washed and re-suspended in PBS. The bacterial inoculum was estimated based on OD_600_ and verified by plating serial dilutions on agar plates to determine colony forming units (CFU). HIOs were cultured in group of 5 per well using 4-well plates (ThermoFisher). Lumens of individual HIOs were microinjected with glass caliber needles with 1 μl of PBS or different strains of *Salmonella* (105 CFU/HIO or 103 CFU/HIO for 24h time point experiments). HIOs were then washed with PBS and incubated for 2hrs at 37°C in ENR media. After 2h, HIOs were treated with 100 μg/ml gentamicin for 15 min to kill any bacteria outside the HIOs, then incubated in a fresh medium containing 10 μg/ml gentamicin.

### ELISA and Bacterial Burden Analyses

Media from each well (5 HIOs/well) were collected at indicated time points after microinjection. Cytokines, chemokines and defensins were quantified by ELISA at the University of Michigan Cancer Center Immunology Core. Bacterial burden was assessed per HIO. Individual HIOs were removed from the Matrigel, washed with PBS and homogenized in PBS. Total CFU/HIO were enumerated by serial dilution and plating on LB agar. To assess intracellular bacterial burden, HIOs were cut open, treated with 100 μg/ml gentamicin for 10 min to kill luminal bacteria, washed with PBS, homogenized and plated on agar plates for 24 hours.

### Immunohistochemistry and Immunofluorescence Staining

HIOs were fixed with either 10% neutral formalin or Carnoy’s solution for 2 days and embedded in paraffin. 5 μm sections were collected by the University of Michigan Cancer Center Histology Core and stained with hematoxylin and eosin (H&E) staining. Carnoy’s-fixed HIO sections were stained with periodic acid-Schiff (PAS) staining kit according to the manufacturer’s instructions (Newcomersupply). H&E and PAS stained slides were imaged on Olympus BX60 upright microscope. All images were further processed using ImageJ.

### RNA Sequencing and Analysis

Total RNA was isolated from groups of 5 HIOs per replicate with a total of 4 replicates per infection condition using the mirVana miRNA Isolation Kit (ThermoFisher). Cytosolic and mitochondrial ribosomal RNAs were removed from samples using the Ribo-Zero Gold Kit according to manufacturer’s protocol (Illumina). RNA samples were used to prepare cDNA libraries by the University of Michigan DNA Sequencing Core and the quality of RNA was confirmed, RIN > 8.5, using the Agilent TapeStation system. Libraries were sequenced on Illumina HiSeq 2500 platforms (single-end, 50 bp read length).

### Reactive Oxygen Species (ROS) Measurement

HIOs were re-plated onto glass-bottom petri dishes (MatTek) and microinjected with 1 μl of PBS/bacteria containing 50 ng/HIO of CM-H2DCFDA (ThermoFisher). HIOs were imaged using inverted widefield live fluorescent microscopy at indicated time points. Images were analyzed by ImageJ.

### Quantification and Statistical Methods

Data were analyzed using Graphpad Prism 7 and R software. Statistical differences were determined using one-way ANOVA or two-way ANOVA (for grouped analyses) and followed-up by Tukey’s multiple comparisons test. The mean of at least 3 independent experiments were presented with error bars showing standard deviation (SD). *P* values of less than 0.05 were considered significant and designated by: \**P* < 0.05, \*\**P* < 0.01, \*\*\**P* < 0.001 and **** *P* < 0.0001.

### Data and Software Availability

Raw data are available upon request, which should be directed to the corresponding authors. Code for analyses can be found at: https://github.com/rberger997/HIO_dualseq2 and https://github.com/aelawren/Salmonella-serovars-RNA-seq.

### RNA-seq analysis protocol

#### Sequence alignment

Sequencing generated FASTQ files of transcript reads were pseudoaligned to the human genome (GRCh38.p12) using kallisto software [24]. Transcripts were converted to estimated gene counts using the tximport package [25] with gene annotation from Ensembl [26].

#### Differential gene expression

Differential expression analysis was performed using the DESeq2 package [27] with *P* values calculated by the Wald test and adjusted *P* values calculated using the Benjamani & Hochberg method [28].

#### Pathway enrichment analysis

Pathway analysis was performed using the Reactome pathway database and pathway enrichment analysis in R using the ReactomePA software package [29].

#### Statistical analysis

Analysis was done using RStudio version 1.1.453. Plots were generated using ggplot2 [30] with data manipulation done using dplyr [31]. Euler diagrams of gene changes were generated using the Eulerr package [32].

## Acknowledgments

This work was supported by the NIAID U19AI116482-01 grant. The funders had no role in study design, data collection and analysis, decision to publish, or preparation of the manuscript. We thank the Host Microbiome Initiative, the Center for Live Cell Imaging (CLCI), Microscopy and Image Analysis Laboratory (MIL), the Comprehensive Cancer Center Immunology and Histology Cores and the DNA Sequencing Core at University of Michigan Medical School. We gratefully acknowledge the O’Riordan lab members for helpful discussions.

## Supporting Information

**Fig S1.**
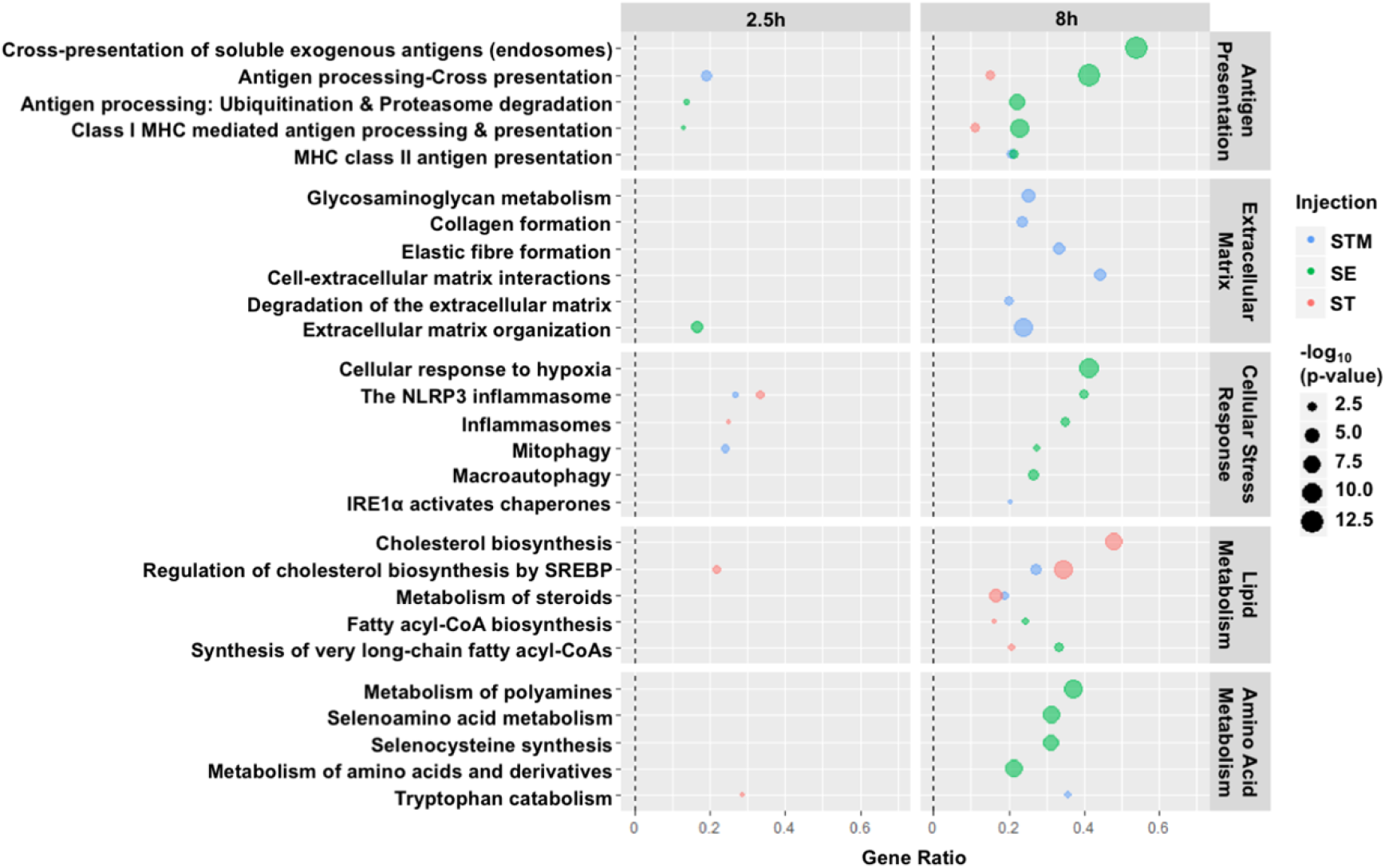
Select Reactome pathways that are differentially upregulated during HIO infection with different *Salmonella* serovars are related to antigen presentation, extracellular matrix, cellular stress responses, lipid metabolism and amino acid metabolism. Dotplot shows select Reactome pathways that are significantly enriched (pValue < 0.05) from upregulated gene sets of HIOs infected with different *Salmonella* serovars relative to PBS-injected HIOs.

**Fig S2.**
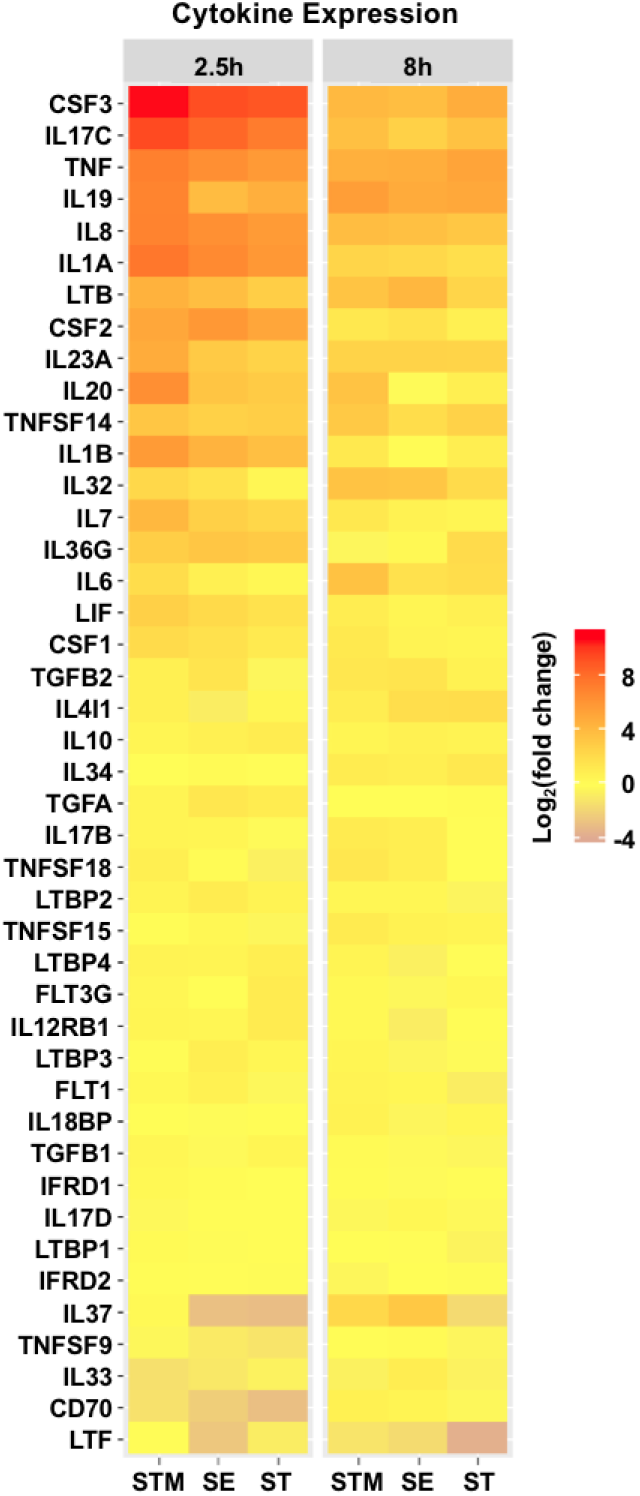
Complete cytokine gene list from Reactome. Gene expression presented as log_2_ fold change relative to PBS injected HIOs at 2.5h and 8h post injection.

**Fig S3.**
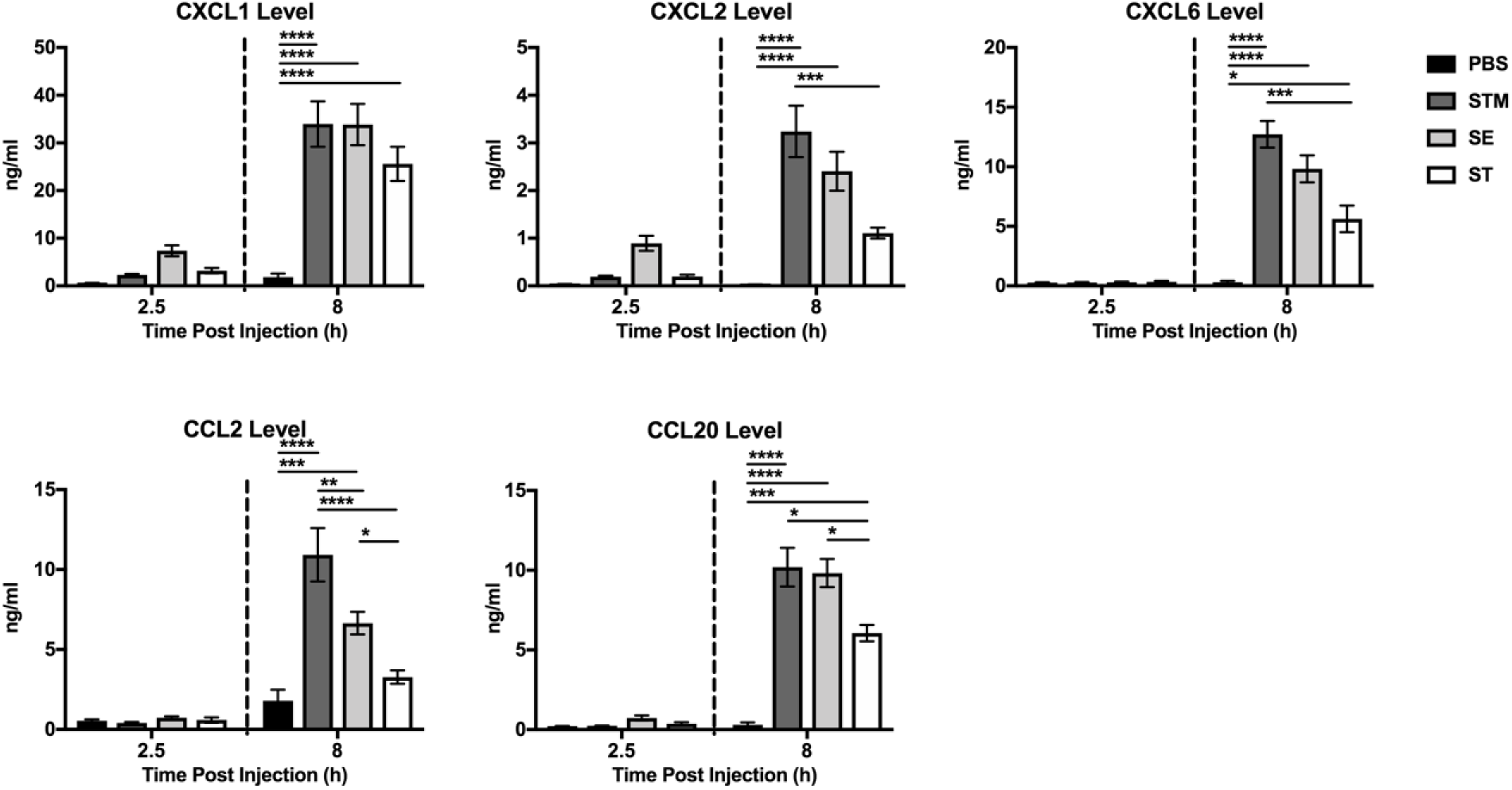
Chemokine secretion levels at 2.5h and 8h post injection for HIOs injected with PBS, STM, SE, and ST. Graphs are presented as mean of n = 4 biological replicates with standard deviation (SD) error bars. *P* value was calculated using two-way ANOVA with Tukey’s post-test for multiple comparisons. \**P* < 0.05; \*\**P* < 0.01, \*\*\**P* < 0.001 and **** *P* < 0.0001.

**S1 Table. DEGs of STM, SE and ST-infected HIOs at 2.5h relative to PBS-injected HIOs.** Significant DEGs (*P* < 0.05) in at least one infection condition are listed.

**S2 Table. DEGs of STM, SE and ST-infected HIOs at 8h relative to PBS-injected HIOs.** Significant DEGs (*P* < 0.05) in at least one infection condition are listed.

**S3 Table. Enriched Reactome pathways from upregulated DEGs at 2.5h pi.** Significantly upregulated DEGs from STM, SE or ST-infected HIOs at 2.5h were subjected to Reactome pathway analysis.

**S4 Table. Enriched Reactome pathways from downregulated DEGs at 2.5h pi.** Significantly downregulated DEGs from STM, SE or ST-infected HIOs at 2.5h were subjected to Reactome pathway analysis.

**S5 Table. Enriched Reactome pathways from upregulated DEGs at 8h pi.** Significantly upregulated DEGs from STM, SE or ST-infected HIOs at 2.5h were subjected to Reactome pathway analysis.

**S6 Table. Enriched Reactome pathways from downregulated DEGs at 8h pi.** Significantly downregulated DEGs from STM, SE or ST-infected HIOs at 2.5h were subjected to Reactome pathway analysis.

